# A Sequence-Pair-Classification-Based Method for Detecting and Correcting Under-Clustered Gene Families

**DOI:** 10.1101/2020.02.22.942557

**Authors:** Akshay Yadav, David Fernández-Baca, Steven B. Cannon

## Abstract

Gene families are groups of genes that have descended from a common ancestral gene present in the species under study. Current, widely used gene family building algorithms can produce family clusters that may be fragmented or missing true family sequences (under-clustering). Here we present a classification method based on sequence pairs that, first, inspects given families for under-clustering and then predicts the missing sequences for the families using family-specific alignment score cutoffs. We have tested this method on a set of curated, gold-standard (“true”) families from the Yeast Gene Order Browser (YGOB) database, including 20 yeast species, as well as a test set of intentionally under-clustered (“deficient”) families derived from the YGOB families. For 83% of the modified yeast families, our pair-classification method was able to reliably detect under-clustering in “deficient” families that were missing 20% of sequences relative to the full/” true” families. We also attempted to predict back the missing sequences using the family-specific alignment score cutoffs obtained during the detection phase. In the case of “pure” under-clustered families (under-clustered families with no “wrong”/unrelated sequences), for 78% of families the prediction precision and recall was ≥0.75, with mean precision = 0.928 and mean recall = 0.859. For “impure” under-clustered families, (under-clustered families containing closest sequences from outside the family, in addition to missing true family sequences), the prediction precision and recall was ≥0.75 for 63% of families with mean precision = 0.790 and mean recall = 0.869. To check if our method can detect and correct incomplete families obtained using existing family building methods, we attempted to correct 374 under-clustered yeast families produced using the OrthoFinder tool. We were able to predict missing sequences for at least 19 yeast families with mean precision of 0.9 and mean recall of 0.65. We also analyzed 14,663 legume families built using the OrthoFinder program, with 14 legume species. We were able to identify 1,665 OrthoFinder families that were missing one or more sequences - sequences which were previously un-clustered or clustered into unusually small families. Further, using a simple merging strategy, we were able to merge 2,216 small families into 933 under-clustered families using the predicted missing sequences. Out of the 933 merged families, we could confirm correct mergings in at least 534 families using the maximum-likelihood phylogenies of the merged families. We also provide recommendations on different types of family-specific alignment score cutoffs that can be used for predicting the missing sequences based on the “purity” of under-clustered families and the chosen precision and recall for prediction. Finally, we provide the containerized version of the pair-classification method that can be applied on any given set of gene families.

## Introduction

Gene families, also known as orthologous groups, are groups of genes from a given set of species that have diverged from one another, from an ancestral gene in the most recent common ancestor of the species. Gene families may contain genes that have diverged due to speciation and/or duplication. Accordingly, genes within a family may be classified as orthologs (separated by speciation) or in-paralogs (duplicated after the common ancestral node) [1–3]. For a number of analysis purposes -- for example, identification of candidates for drug/vaccine development [4–6] or annotation of newly sequenced genomes by cross-referencing function information from multiple species [7–11] -- it is useful to identify families such that all member genes originated from a single ancestral gene that was present in the common ancestor of the species under study. Many popular clustering techniques use these basic evolutionary properties of gene families for building gene families from whole proteomes of the species. These clustering algorithms use some form of normalized similarity/alignment scores between sequences as an input to a clustering method such as Markov Clustering (MCL) [12, 13] to generate gene family clusters. Two of the most popular family building methods, OrthoFinder [14] and OrthoMCL [15], use normalized BLAST [16, 17] scores or E-values to cluster sequences into families using the chosen clustering algorithm. For Markov Clustering, which is widely used, the granularity of the MCL clusters is controlled by the inflation (I) parameter, with higher values of the parameter generally corresponding with a larger number of clusters. Both OrthoFinder and OrthoMCL use a single value of the Inflation parameter for building all families for a given clustering run. Since different gene families can evolve at different rates [18–21], using a single Inflation parameter value for MCL clustering may be over-stringent for some families (resulting in fragmented/under-clustered families) - and under-stringent for other families (resulting in merged/over-clustered families).

Here, we present a classification method based on machine learning that first inspects a given family for under-clustering in a training step, and subsequently attempts to predict missing sequences for the family using family-specific alignment score cutoffs obtained in the training step. The training step consists of repetitive model building and testing where, during each iteration, a combined set of sequences from the given family along with a selected set of closest non-family sequences is randomly split into training and testing parts. The training part of the family is used to build a HMM [22], which is then tested to recognize the correct family sequences from the testing part, containing both family and non-family sequences. Pairs of sequences were used as data points for training and testing the family models. Using pairs helps to create more data for training and testing, giving ^*N*^*C*_*2*_ pairs (data points) for training and testing, from a family with N sequences.

## Methods

### Collecting Candidate Missing Sequences

Sequences from a given family were searched against the database of proteomes from all the species under study. With each family sequence as query, all the hits that match the query better than the worst-matching family hit were collected into a list. Each query sequence can attract one or more non-family sequences along with the original family sequences. A combined list of sequences was prepared from the lists of hits collected from searching all the family sequences against the database (Fig 2.1). This combined list contains all of the original family sequences and can contain one or more non-family sequences. The non-family sequences in the list of retrieved sequences are candidates for sequences missing from the family.

**Fig 2.1.**
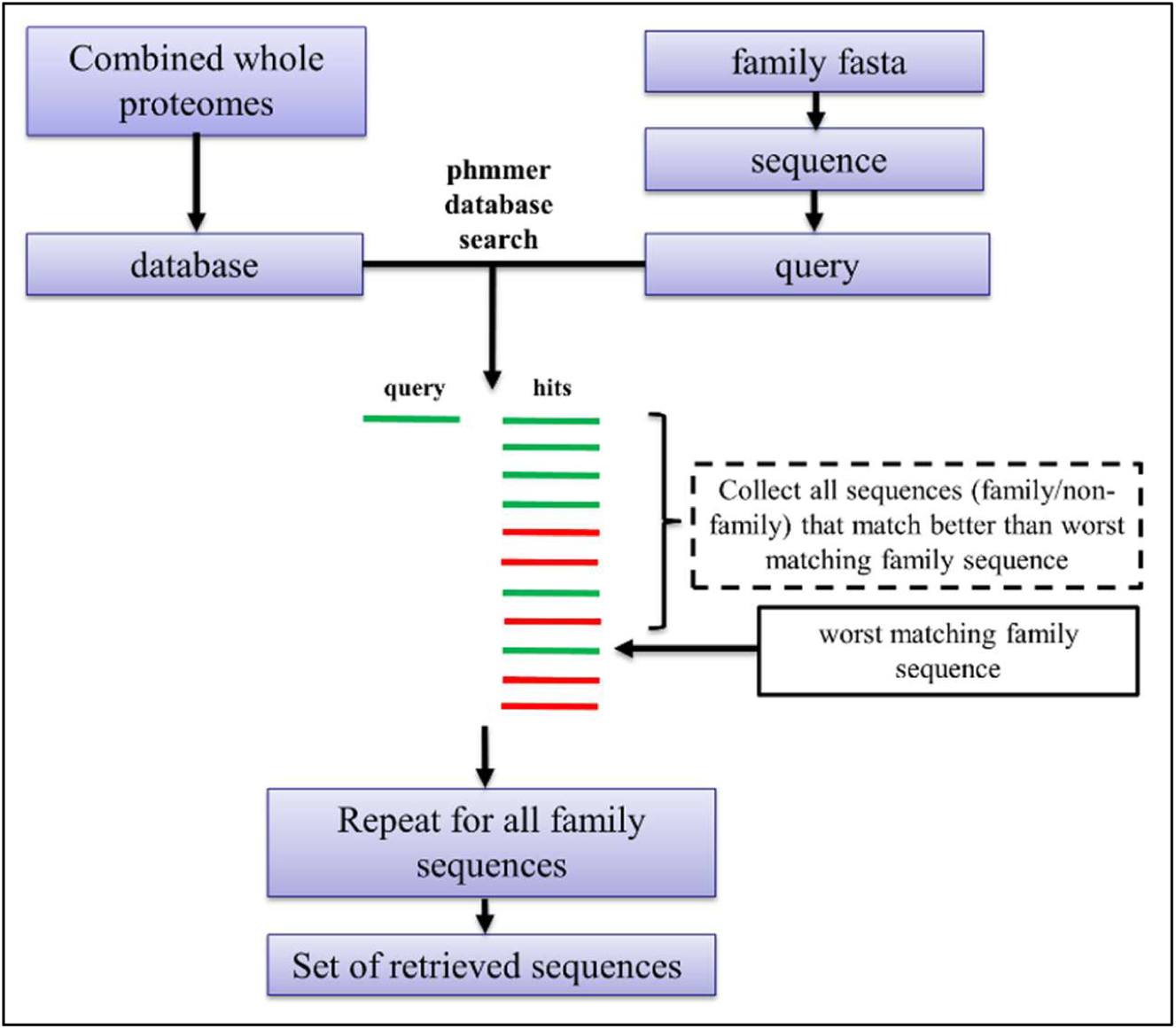
Procedure for collecting candidate missing (non-family) sequences for a given family. Each sequence from a family was searched against the proteomes from target and outgroup species using the phmmer program. For every query sequence, all the hits that match the query with better scores than the worst-matching family sequence were added to a set of retrieved sequences containing all the original family sequences, plus the closest non-family sequences. The non-family sequences are candidates for sequences missing from the original family.

The phmmer program from the HMMER package (version 3.1b2) [23] was used for searching families against the database of whole proteomes. E-values from phmmer output were used to rank the hits, in order to find the worst-matching family hit.

### Defining Sequence Pairs

The list of retrieved sequences obtained in the previous step was used to define family and non-family sequence pairs (Fig 2.2). Family pairs are those exclusively between original family members, and non-family pairs are those between family and non-family members. Family pairs were labeled as “positive” and non-family pairs were labeled as “negative”.

**Fig 2.2.**
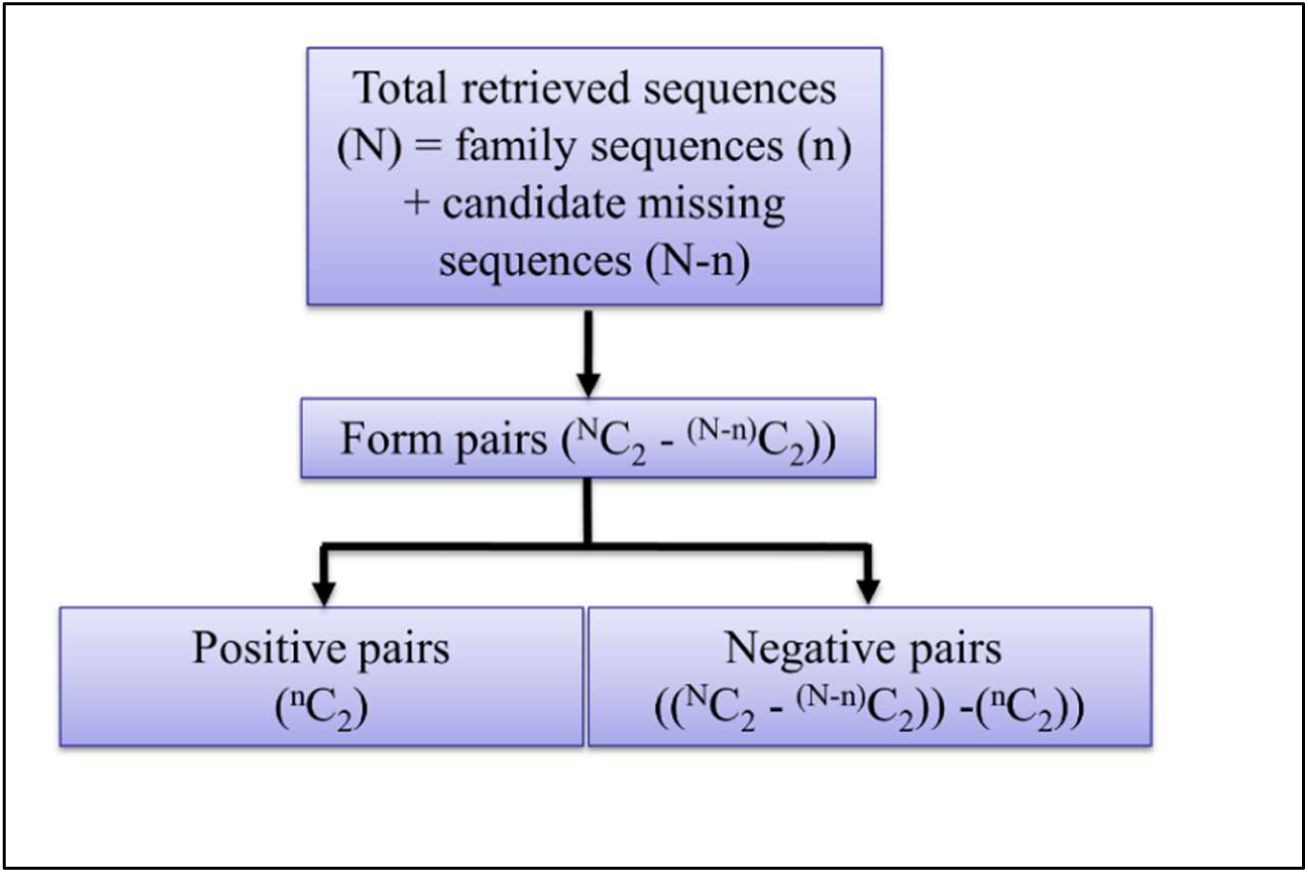
Defining positive and negative pairs from the family and non-family sequences obtained in the previous step. Let ‘N’ be the total number of sequences in the list of retrieved sequences for a given family and let ‘n’ be the number of original family sequences. Then, (N-n) is the number of non-family sequences in the retrieved list of sequences that could be missing from the family. Pairs are only formed between the family members (positive pairs) and between the family members and non-family sequences (negative pairs). The number of pairs that can be formed from a set of retrieved sequences containing ‘N’ sequences with ‘n’ sequences from the original family is (^*N*^*C*_*2*_ *-* ^*(N-n)*^*C*_*2*_). From these pairs, there are ^*n*^*C*_*2*_ positive pairs and ((^*N*^*C*_*2*_ *-* ^*(N-n)*^*C*_*2*_*)-* ^*n*^*C*_*2*_) negative pairs.

### Training and Classification Statistics

HMM-based pair-classification models were built to classify the positive (family) pairs from negative (non-family) pairs, for a given family, and 10 iterations of repeated test/train split strategy were used to assess the classification performance (Fig 2.3). For each iteration, the set of positive pairs was randomly split into training (80%) and test (20%) sets. The HMM model was trained on consensus sequences of positive pairs from the training split, obtained using the “-c” option of hmmemit program (HMMER package:version 3.1b2). The MAFFT program (version v7.407) [24] was used for calculating the multiple sequence alignment used for building the HMMs. Subsequently, the trained HMM was tested on unseen positive pairs in the test split and the negative pairs by aligning individual sequences of the pairs to the HMM using the hmmsearch program. For each test pair, full sequence alignment scores for both the sequences were obtained. The test pair was predicted as positive if both the alignment scores were greater than or equal to a specified alignment score cutoff, else the test pair was predicted as negative.

**Fig 2.3.**
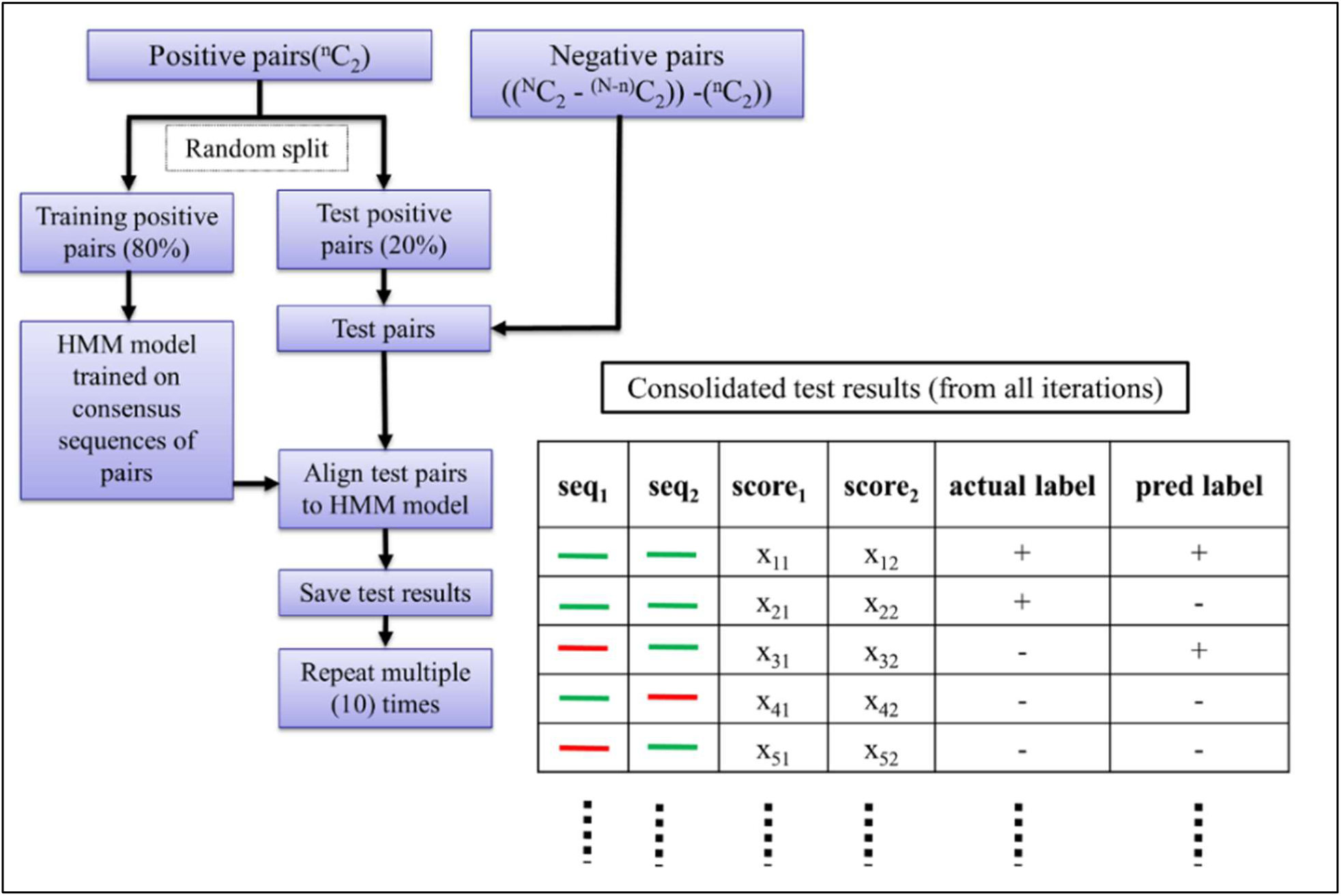
Model training and testing for classifying the positive pairs from the negative pairs. For each iteration, the set of positive pairs was randomly split into training and testing sets, with the training set containing 80% of positive pairs and the testing set containing the remaining 20%. Consensus sequences of positive pairs from the training set were used to build HMM and the positive pairs from the test set and all the negative pairs, were aligned to the HMM. Individual sequences of each test pair were aligned to the HMM and full sequence alignment scores for both sequences were recorded. Alignment score results obtained from aligning unseen test pairs to trained HMMs, consolidated from all test-train split iterations performed on a given family, were used for calculating classification statistics for the family to analyze the separation between positive and negative pairs. The alignment scores of the individual sequences of the test pairs were used to predict the test pairs as positive or negative. Given a fixed score cutoff, a test pair was predicted as positive if both alignment scores corresponding to both the sequences of the pair were greater than or equal to a score cutoff, else the test pair was predicted as negative. Accordingly, for a fixed score cutoff, True Positive (TP) test pairs were those that were originally positive and were also predicted as positive, False Positive (FP) test pairs were those that were originally negative but were predicted as positive, and finally, False Negative (FN) test pairs were those that were originally positive but were predicted as negative.

The alignment scores for test pairs from all the iterations were consolidated (Fig 2.3) and used for calculating precision (TP/TP+FP), recall (TP/TP+FN) and F-score (eq 1) values for specified alignment score cutoffs, where TP, FP and FN are the number True Positive, False Positive and False Negative pairs, respectively. The F-score is defined as

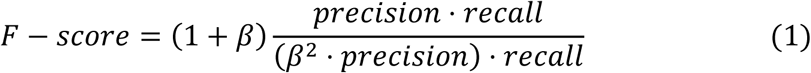

where β controls the importance of importance of recall over precision with values greater than 1 favoring recall over precision [25]

Precision and recall values were obtained for a range of alignment score cutoffs, ranging from most to least stringent. Precision values were plotted against the corresponding recall values to obtain the Precision-Recall curve (PR-curve) [26] and the area under the PR-curve (PR-AUC) was calculated using the trapezoidal rule [27]. The PR-AUC value ranges from 0 to 1 and was used as a score indicating family completeness. Complete families with no missing sequences are expected to have PR-AUC values closer to 1, indicating good separation between the positive and negative pairs. An example of PR-curve plot for a hypothetical gene family is shown in Fig 2.4. Each point on the curve corresponds to a recall value and the corresponding precision value (recall, precision) obtained using an alignment score cutoff with score cutoffs decreasing from left to right. The score cutoffs on the left produce pair classifications with high precision (low FP) but low recall (high FN). Conversely, low score cutoffs on the right produce pair classifications with low precision but high recall. An F-score can be calculated for each point (recall, precision) on the curve using the F-score function (eq 1). The point on the curve with the highest F-score is the point where optimal values of precision and recall exist (optimal trade-off between precision and recall). This point represents the best classification performance for the family and the corresponding score cutoff gives the best possible separation between the positive and negative pairs of the respective family.

**Fig 2.4.**
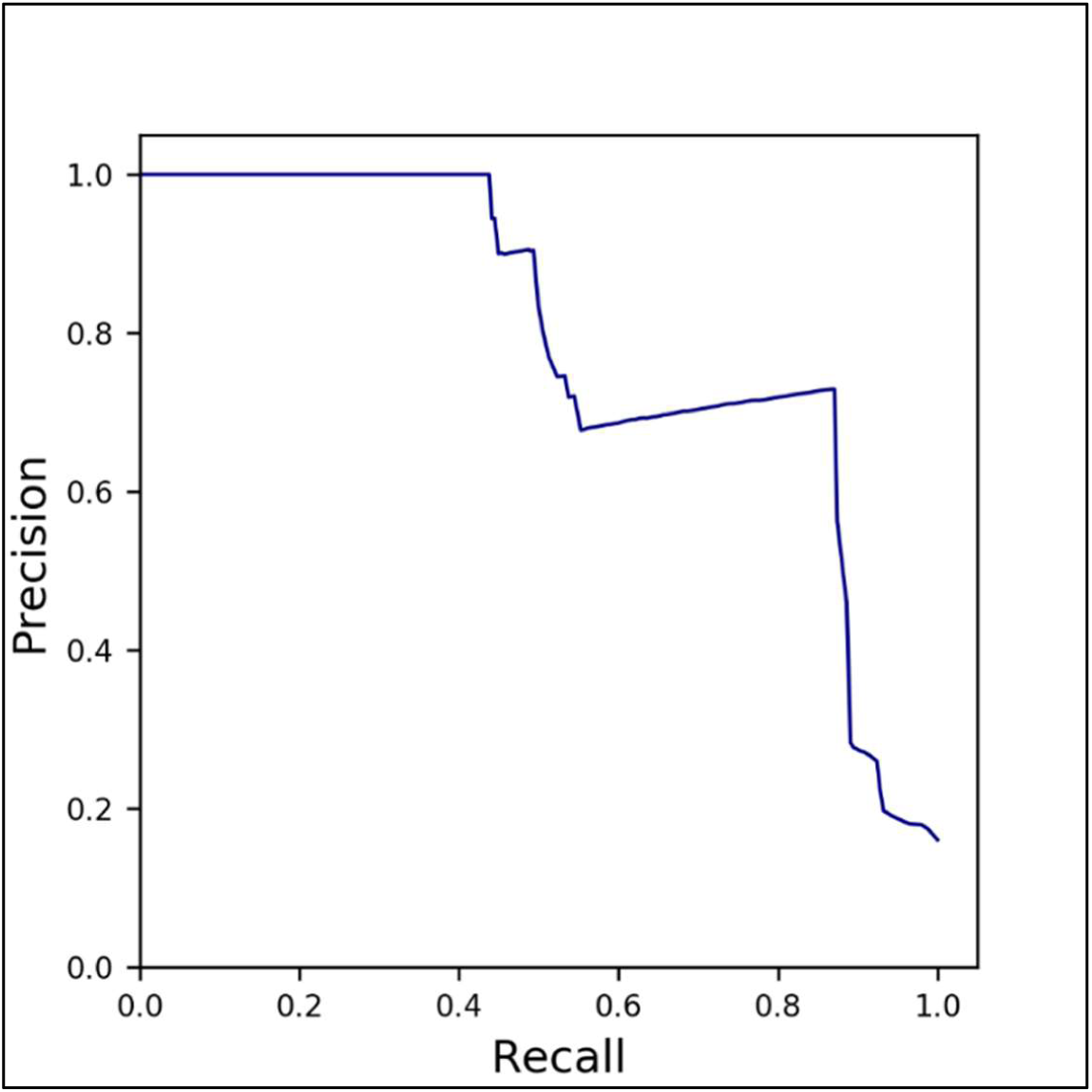
A typical Precision-Recall (PR) curve for a gene family. The precision and recall values obtained using different alignment score cutoffs were plotted against each other to obtain the Precision-Recall curve (PR-curve). The alignment score cutoffs vary from high value to low value from left to right, with cutoffs on the left giving high precision but low recall and cutoffs on the left giving low precision and high recall, for classification between the positive and negative pairs of a given family. The area under the PR-curve (PR-AUC) was calculated using the trapezoidal rule.

Classification metrics such as the precision and recall values observed at the best F-score and the alignment score cutoff which gives the best F-score were calculated. Two types of alignment score cutoffs were reported corresponding to the two values of the β parameter (β = 1 and β = 2) in the F-score function. The F-score function with β = 1 is called the F1-score function and the F-score function with β = 2 is called the F2-score function. In addition, the lowest alignment score observed for the positive pairs was also reported as the lowest alignment score cutoff for the given family.

### Predicting Missing Sequences

Missing sequences for every family were predicted using the negative pairs (Fig 2.5). A single HMM was built using the consensus sequences of all the positive pairs. Subsequently, all the negative pairs were aligned to this HMM and those negative pairs where the alignment scores for both the sequences of the pairs were greater than or equal to the chosen type of score cutoff (F1/F2/lowest) were re-classified as positive pairs. Unique sequences within these re-classified positive pairs were predicted and reported as the missing sequences for the family. The precision and recall values for prediction of missing sequences were also calculated, as (TP/(TP+FP)) and (TP/(TP+FN)), respectively, where True Positive (TP) are those predicted missing sequences that were truly missing from the family, False Positive (FP) are those that were predicted as missing but do not actually belong to the family and False Negative (FN) are those that are truly missing but were not predicted as missing.

**Fig 2.5.**
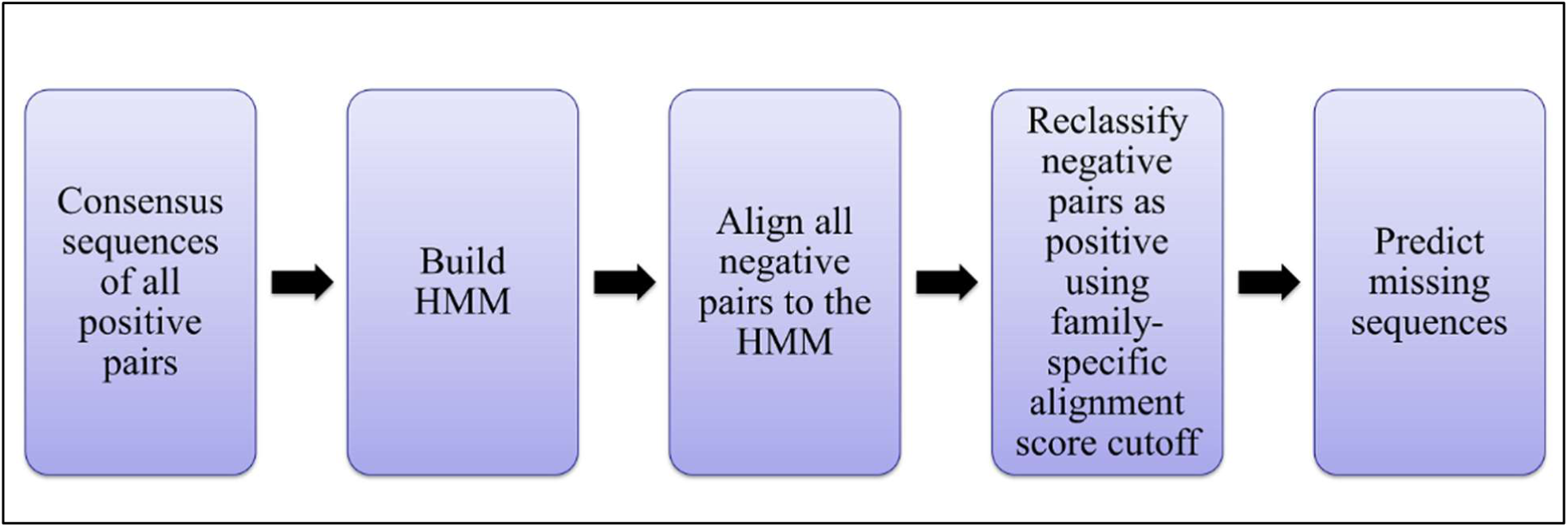
Predicting missing sequences using the negative pairs and family-specific alignment score cutoff. For each family, consensus sequences of the positive pairs and the negative pairs were aligned to the trained HMM for the family. Negative pairs where alignment scores of both the sequences of the pair were greater than or equal to the family-specific alignment score cutoff were reclassified as positive pairs, and the constituent sequences were predicted as missing sequences.

## Results

### Behavior on “true” YGOB Families

The pair-classification-based scoring was tested on 4,796 yeast families from the Yeast Gene Order Browser (YGOB) database [28]. Since the YGOB families are built through manual curation using synteny-based evidence, they were assumed to be correct or “true.” To check if the pair-classification method is assigning high classification performance scores to all the true families, the distribution of the PR-AUC values for all the 4,796 yeast families was obtained (Fig 2.6A). As expected, the distribution is highly skewed towards PR-AUC value = 1.0, with 92% of family classifiers having values ≥0.75. This shows that the proposed method correctly recognizes complete families and assigns high classification performance scores to them.

**Fig 2.6.**
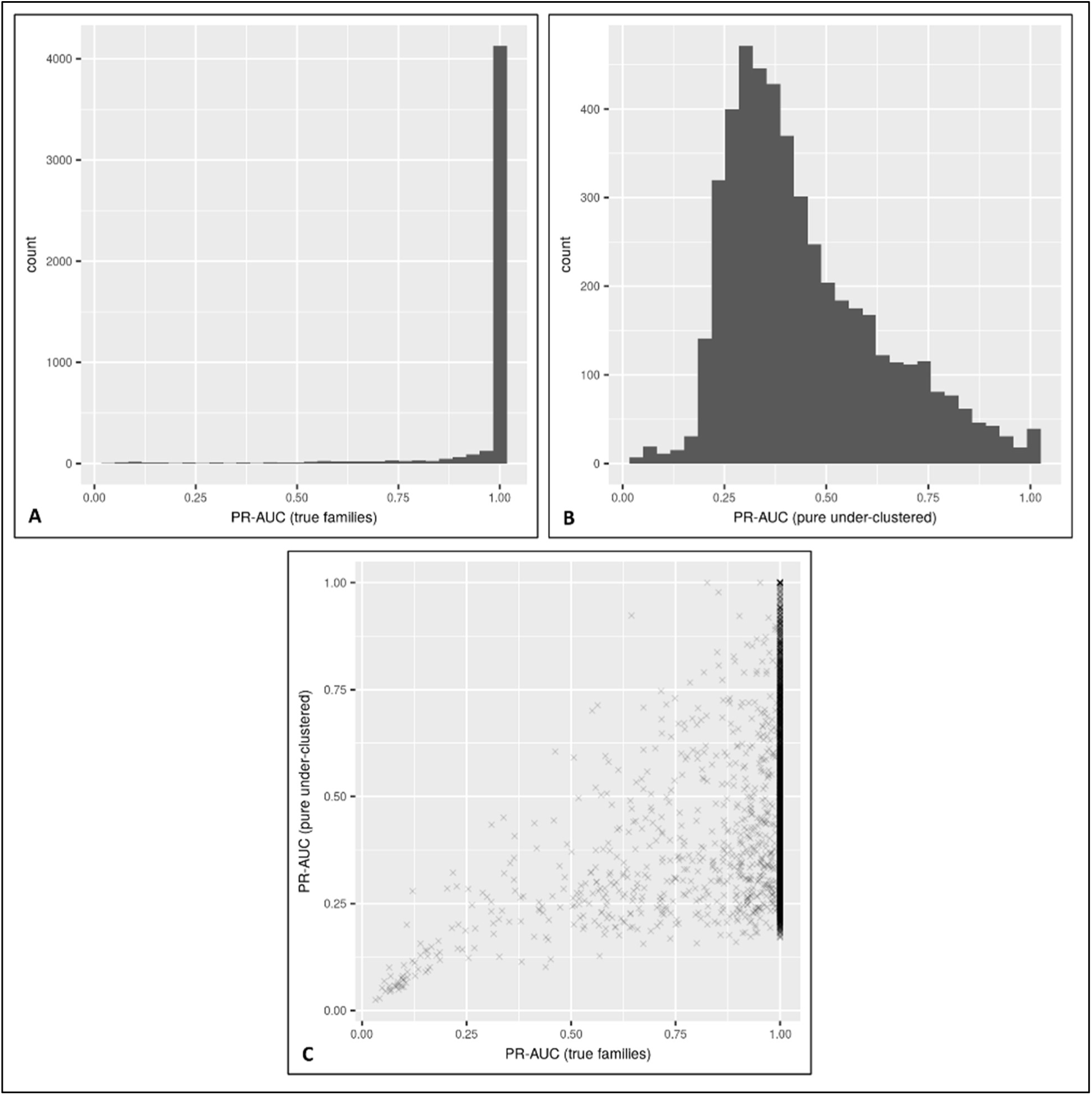
(A) Distribution of PR-AUC values for “true” yeast families. (B) Distribution of PR-AUC values for “pure under-clustered” yeast families with 20% of family sequences removed. (C) Scatterplot comparing the PR-AUC values of true families vs. incomplete families with 20% of sequences deleted. The distribution for true families is heavily skewed towards 1.0 which shows that separation between the positive and negative pairs is good and in majority of the cases perfect, for true gene families. The perfect classification signifies that the families are complete with no missing sequences. The distribution for pure under-clustered families shows the drop in the PR-AUC values as compared to the distribution of PR-AUC values for true families. The scatterplot shows the drop in the PR-AUC value for each family after 20% of the sequences are removed from the family. Each point in the plot represents a family.

### Results for correction of “pure” but under-clustered families

Even though this under-clustering assessment method performs well in our tests on good/true families, it is important to study the behavior of the method on incomplete families. To check the behavior of the method on incomplete families, the “true” yeast families were modified so that each family is missing a random 20% of the family sequences.

The distribution of PR-AUC values for 4,796 artificially manipulated families where every family is missing 20% of their sequences is shown in Fig 2.6B. The distribution indicates that the pair-classification performance of the families drops significantly, which shows the ability of proposed method to detect incomplete families. For 3,971 out of 4,796 families (83%), the PR-AUC value dropped significantly: PR-AUC ≥ 0.9 for true families and PR-AUC < 0.75 after removing family sequences (Fig 2.6C).

For each pure under-clustered family, the missing sequences were predicted back using lowest alignment score cutoff obtained during training. Since these incomplete families are “pure” i.e. they do not contain any non-family sequences, the lowest score cutoff can be regarded as a lower bound of the family. Any true family sequence/sequence pair is expected to align to the family HMM with a score greater than the lowest score cutoff. The predicted missing sequences were compared to the true missing sequences, for each family, and precision and recall values were calculated to study the accuracy of prediction. Out of 4,796 families, the prediction performance for missing sequences was high (precision ≥ 0.75 and recall ≥ 0.75) for 3760 (78.4%) families with overall mean precision = 0.928 and overall mean recall = 0.859.

### Results for Correction of “impure” and Under-clustered Families

Under-clustered families can also contain non-family or “wrong” sequences. To evaluate the behavior of our method on incomplete families contaminated with unrelated sequences, 2,391 yeast families were modified so that each family contained an additional 20% of sequences, from a set of the closest non-family sequences - in addition to missing 20% of the original family sequences. These were called “impure, under-clustered families” and were analyzed using the pair-classification method. Since these under-clustered families contain unrelated sequences, it is possible that more unrelated sequences can be attracted while selecting the candidate missing sequences, in the first step of the workflow (See methods). To make sure only relevant missing candidates are selected, only those non-family sequences were retained in the first step that were attracted by at least 50% of family sequences.

As observed in the case of pure under-clustered families, there was a significant reduction in the PR-AUC values for 83% of impure, under-clustered families, as compared to the true families, indicating that the pair-classification method is able to detect under-clustering even when there are wrong sequences present in the under-clustered family. Since these families already contain wrong/unrelated sequences, the lowest score cutoff that gives highest recall cannot be used for predicting missing sequences. Therefore, two different types of alignment score cutoffs - score cutoff obtained using the F1-score function and score cutoff obtained using the F2-score function, were used to predict the missing sequences for each of the 2,391 impure under-clustered families. Table 2.1 shows the precision and recall results for prediction of missing sequences obtained using the two types of score cutoffs.

**Table 2.1.** Precision and recall results for predicting missing sequences, for yeast families containing non-family sequences in addition to missing 20% of the sequences

| **Function type** | **% of families with prediction precision $\geq 0.75$** | **% of families with prediction recall $\geq 0.75$** | **% of families with both prediction precision $\geq 0.75$ and prediction recall $\geq 0.75$** | **mean prediction precision** | **mean prediction recall** |
| --- | --- | --- | --- | --- | --- |
| F1-score | 1729/2391 = 72.3 | 1348/2391 = 56.3 | 1138/2391 = 47.5 | 0.766 | 0.666 |
| F2-score | 1685/2391 = 70.4 | 1978/2391 = 82.7 | 1426/2391 = 63.1 | 0.790 | 0.868 |

The prediction results show that the alignment score cutoffs obtained using the F1-score function predicts missing sequences with high precision for 72% of the families but fails to recognize true missing sequences for a majority of the families, with only 56% of the families having high recall. Consequently, the overall prediction performance is high (high precision with high recall) for only 47% of the families with mean precision = 0.766 and mean recall = 0.666. In order to increase the recall performance, score cutoffs obtained using F2-score were also used to predict missing sequences for same set of impure under-clustered families. The F2-score function is expected to give alignment score cutoffs that favor recall over precision. Accordingly, the alignment score cutoffs obtained using the F2-score function improved the recall for prediction of missing sequences with 82% families having high recall and with 70% families having high precision performance with mean precision = 0.790 and mean recall = 0.868. This has also increased the overall prediction performance with 63% of families having high prediction precision and recall for predicting missing sequences.

### Comparison to Existing Family Building Methods

To check if the pair-classification method can improve families built using existing family building methods, we analyzed 374 under-clustered yeast families obtained using the OrthoFinder tool. The OrthoFinder method has been shown to outperform many popular family building methods like OrthoMCL, TreeFam and OMA [15, 29, 30]. The pair-classification method was able to improve 5 to 19 under-clustered families by predicting missing sequences using the family-specific alignment score cutoffs obtained using F1-score and F2-score functions, respectively. Table 2.2 shows the results on improving the under-clustered OrthoFinder families using two types of alignment score cutoffs obtained using the two F-score functions.

**Table 2.2.**
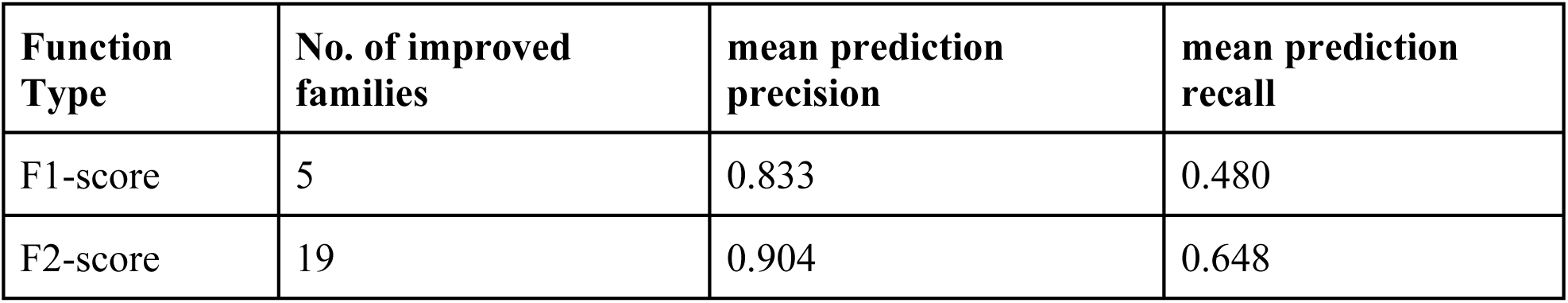
Results on correcting under-clustered yeast families obtained using the OrthoFinder tool.

As expected, the more conservative F1-score function is able to correct only 5 families with high precision of 0.833 and low recall of 0.48. In comparison, the F2-score function is able to improve 19 incomplete families, with significantly more recall of 0.65.

### Analyzing and Correcting OrthoFinder Legume Families

We also analyzed 14,663 legume families built using the OrthoFinder (version 2.2.0) tool from 14 legume proteomes [31–42] (Table 2.2), using the pair-classification method. The 14 legume species belong to subfamily Papilionoideae of family Fabaceae (the third largest family of flowering plants [43]). An ancient Whole Genome Duplication (WGD) occurred in the common ancestor of the Papilionoid sub-family, around 55 Ma [44–49]. In addition, some genera such as *Glycine* and *Lupinus* have also undergone independent, lineage specific WGDs [49, 50].

**Table 2.2.**
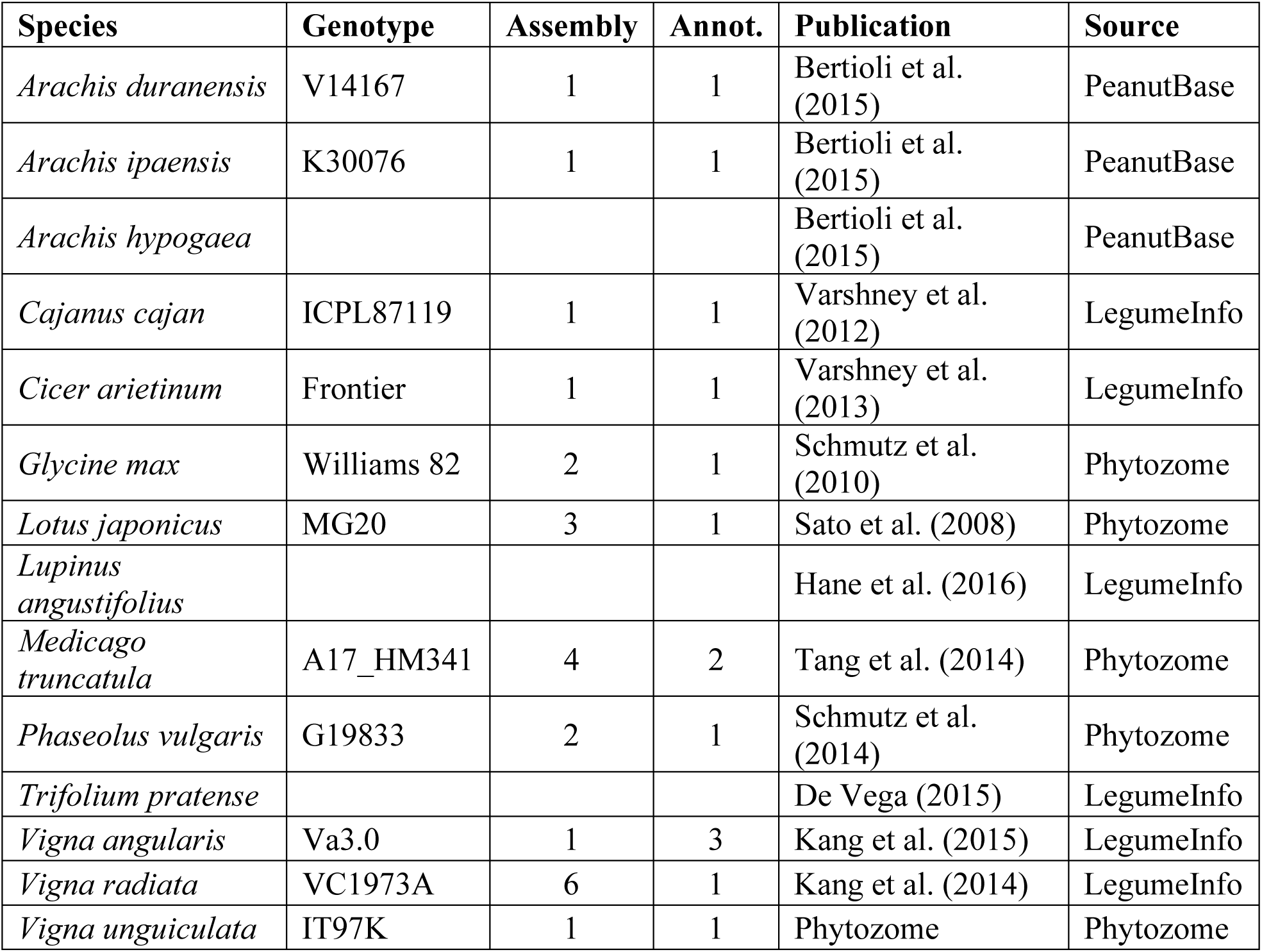
Genome and annotation sources and versions.

The distribution of family sizes up to size 100 is shown in Fig 2.8. Approximately 12% of sequences (64,047) were not clustered by OrthoFinder. Considering families with sizes <=8 as unusually small and assuming the unclustered sequences as singleton families, there were 10,963 + 64,047 = 75,010 unusually small families with sizes ≤ 8 where the 10,963 families were those that contain 2 to 8 sequences.

**Fig 2.8.**
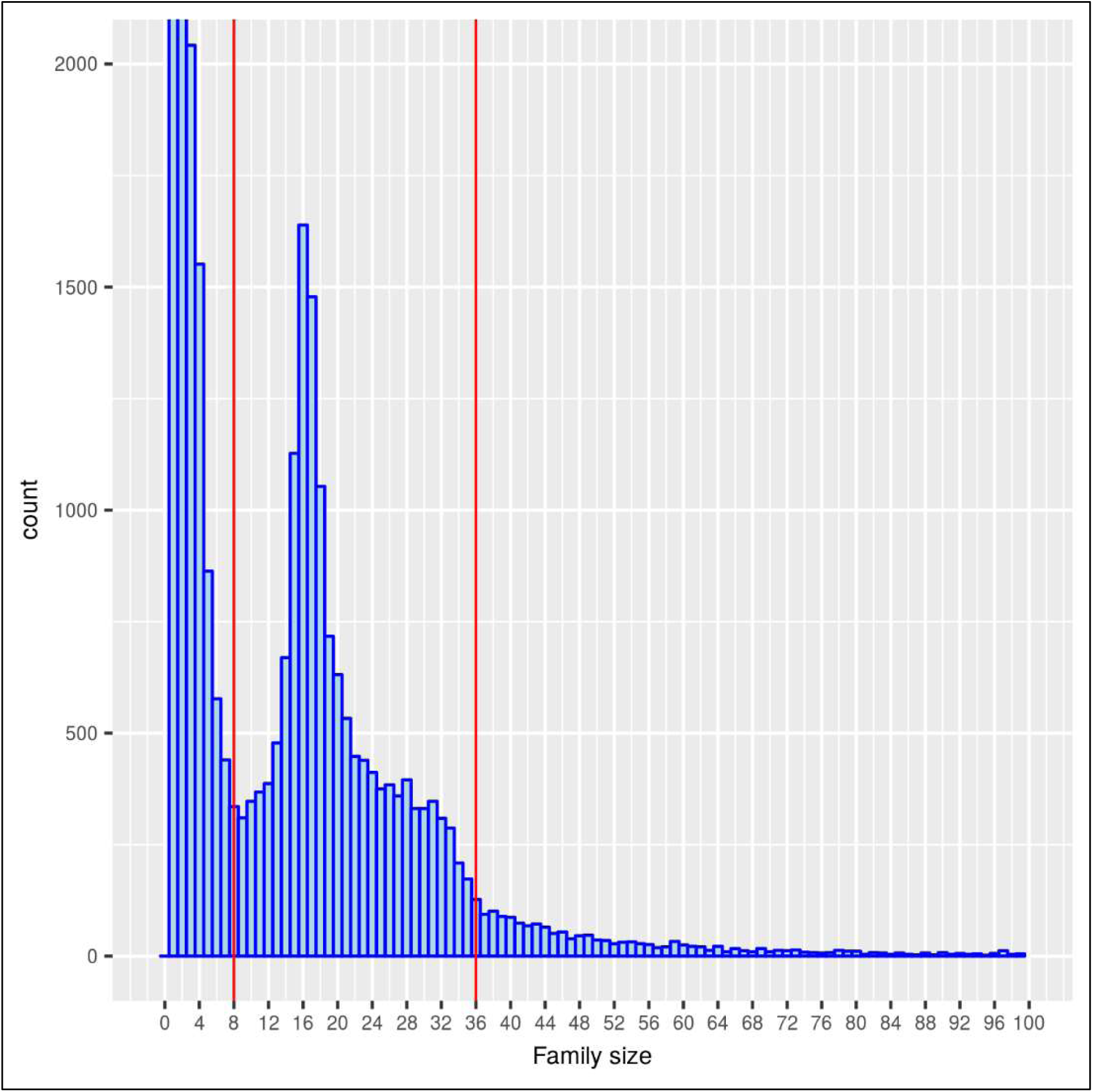
Family size distribution for legume families built using the OrthoFinder tool. The size distribution is shown up to size 100. The families with sizes between 1 and 8 were considered unusually small. The families that fall between the vertical red lines (9 ≤ size ≤ 36) were analyzed using the pair-classification method.

Our hypothesis is that these small families and the unclustered sequences could be produced as a result of over-fragmentation or under-clustering of larger families, due to stringent clustering parameters and can be merged into the larger families. Accordingly, 14,663 larger families with sizes between 9 and 36 were analyzed using the pair-classification method. The unclustered sequences together with sequences from smaller families with sizes between 2 to 8 were compiled together to be used as the sequence database for phmmer search, for gathering candidate missing sequences for each of the 14,663 families, in the first step of the workflow (see methods). Only those non-family sequences were selected as candidate missing sequences that were attracted by at least 50% of family sequences. Out of the 14,663 families, for 9581 families, no candidate missing sequences were found according to this selection criteria.

For the rest of the 5082 families, for which one or more candidate missing sequences were detected, pair-classification models were built and tested to study the separation between the positive pairs formed within the family sequences and the negative pairs formed between the family and non-family sequences. The distribution of PR-AUC values for the 5082 legume families is shown in Fig 2.9. The distribution is skewed towards 1.0 which shows that the majority of these families show good separation between the positive and negative pairs.

**Fig 2.9.**
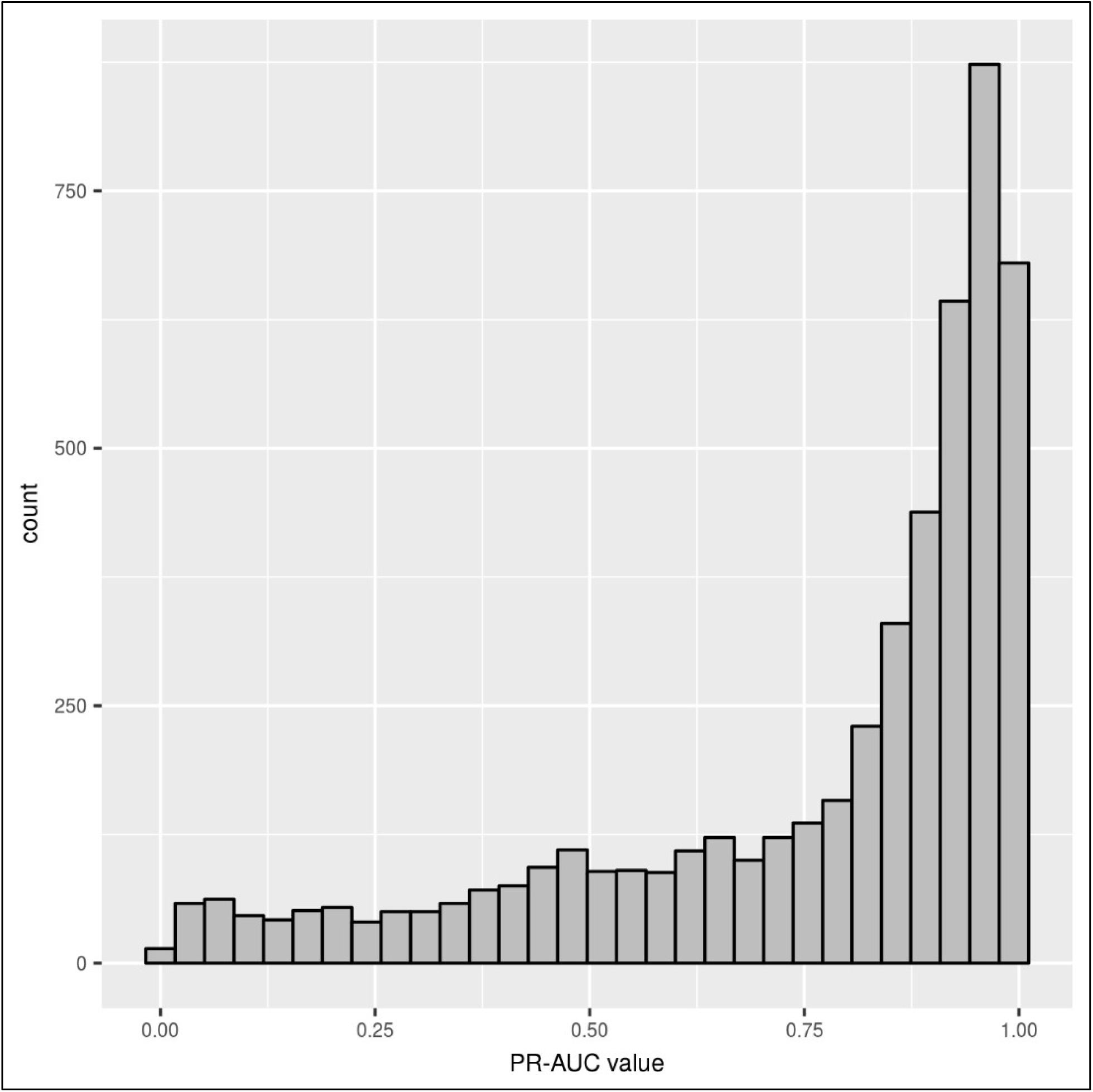
Distribution of PR-AUC values for 5082 families analyzed using the under-clustering detection and correction method. The distribution is skewed towards higher PR-AUC values, signifying good classification performance for the majority of the families.

For 1,665 families out of 5,082, one or more missing sequences were predicted using the family-specific alignment score cutoffs obtained in training, using the F1-score function. The F1-score function was used to obtain the optimal alignment score cutoffs as family precision was considered more important than recall, for predicting the missing sequences. In total, there were 3,588 sequences from the small families that were predicted as missing for these 1665 larger families.

Next, we attempted to merge the smaller families into the larger families using the predicted missing sequences with the following merging rule. If, for a larger family, missing sequences were predicted from a smaller family that were more than 50% of the size of the smaller family, the corresponding smaller family was merged into the larger family. For small families that had potential to merge into more than one larger family, the hhsearch program (version 2.0.16) [51], from the hh-suite package, was used to select the best-matching larger family. In all, 2,216 small families were merged into 933 larger families using the pair-classification method.

We also used a tree-based method to check the correctness of the family mergings. For each of the 933 merged families, closest outgroup sequences with respect to the legumes were identified using the respective family HMMs. The family HMMs were searched against the database of outgroup sequences and up to 10 top-matching outgroup sequences, with e-value ≤1e-5, were selected as the closest outgroup sequences. Phylogenetic trees were calculated for each of the merged families along with their respective outgroup sequences using RAxML [52], with the PROTGAMMAAUTO substitution model, and were rooted using the closest available outgroup species. Each family tree was then analyzed for presence of monophyletic clades containing all the legume sequences. The merging was considered correct if only one monophyletic legume clade was observed in the merged family tree. Out of the 933 trees, 478 families were found to contain single legume monophyletic clade. Also, there were 56 families which contained more than one legume clades but the newly merged sequences were part of a major clade containing 70% or more sequences from the merged family, and the minor clades contained 30% or less sequences that were part of the original unmerged family. For at least 534 families, out of the 933 corrected families, family mergings predicted by the pair-classification method were consistent with expected phylogenetic relationships.

We also attempted to predict missing sequences for the 14,633 legume families using a simple HMM searching strategy, to highlight the importance of family-specific alignment score cutoffs. Every family HMM was searched against the database of sequences from the small families and sequences that align to the family with e-value ≤ 1e^-10^ were predicted as missing sequences for the family. This resulted in unusually large family clusters with more than 1600 clusters containing ≥ 100 sequences, and the largest clusters containing more than 1400 sequences. This showed that the generic E-value cutoff is too relaxed for some families and is attracting a large number of sequences. For example, for family OG0001825 (size = 36), 6 sequences were predicted as missing using the family-specific alignment score cutoff through the pair-classification method, as opposed to 227 sequences that were predicted as missing using the simple HMM searching strategy with a e-value cutoff of 1e^-10^.

## Discussion

Under-clustering is a common problem in current family building methods. For example, at least 374 incomplete yeast families were produced by the OrthoFinder tool. Similarly, in the case of legume families, the OrthoFinder method could not assign about 12% of the genes to any family, and many small families were also produced - potentially indicating fragmentation of larger families. Here, we present a sequence-pair-based classification method that is not only able to detect whether a given family is under-clustered or not but can also predict the missing sequences for the incomplete families.

To check the effectiveness of the method, it was tested on “true” and modified yeast families. On the true yeast families, the method correctly identified complete families, assigning near-optimal or perfect PR-AUC values to the unmodified families, and also identifying under-clustered families through low or sub-optimal values for the PR-AUC statistic. The results on prediction of missing sequences for the modified families showed that the family-specific alignment score cutoffs obtained during training the pair-classification models were able to recognize true missing sequences for the families even when unrelated sequences were present in the families. To check if the pair-classification method can improve families built using existing methods, we applied this method for correcting 374 incomplete family clusters produced by the OrthoFinder method. The pair-classification method was able to improve up to 19 families by predicting the correct missing sequences for these families with mean precision of 0.9 and mean recall of 0.65. Finally, we also applied this method for analyzing 14,633 legume families built using the OrthoFinder method. The pair-classification method was able to identify 3,588 missing sequences for 1,665 families which were subsequently used to merge 2,216 small families into 933 larger families using a simple merging rule. Comparisons against phylogenies of the merged families provided confirmatory evidence for mergings in at least 534 of the 933 merged families.

Different types of family-specific alignment score cutoffs can be used for predicting the missing sequences depending upon the nature of under-clustering and preference of family precision or family completeness. There is a tradeoff between the objectives of family accuracy and family completeness. If family precision is valued more highly than family completeness, then an alignment score cutoff with high value is recommended. Conversely, if family completeness (recall) is preferred over precision, then a low alignment score cutoff should be used for predicting missing sequences. The alignment score cutoff obtained using the F1-score function appears to be predicting missing family sequences with high precision and low recall, as seen from the prediction results on impure under-clustered families. Therefore, the alignment score cutoff obtained using the F1-score function can be used to predict missing sequences for families with high precision. On the other hand, the alignment score cutoffs obtained using the F2-score function improves the recall for prediction at the expense of precision, favoring recall over precision. The high-recall alignment score cutoff obtained using the F2-score function can be used for predicting missing family sequences with more recall at the expense of precision. The lowest alignment score cutoff with the highest possible recall can be used in case of those under-clustered families that are not “contaminated” with unrelated sequences.

Based on the results obtained from modified under-clustered yeast families, we can make the following recommendations on which type of alignment score cutoff to use for a given set of families. If the user/expert is confident that any family detected as under-clustered (has a low PR-AUC value) has a low probability of containing non-family/unrelated sequences, then the score cutoff obtained using the F2-score or the lowest possible score cutoff can be reliably used for predicting the missing family sequences. In contrast, if the user thinks that the under-clustered family may contain up to 20% of unrelated sequences, then the more conservative score cutoff obtained using the F1-score function should be used for predicting the missing sequences.

The method presented here can be used as a post-processing tool for independently assessing gene family sets built using existing family building methods. Every family can be analyzed using the pair-classification method for signs of under-clustering, using the PR-AUC value obtained during training the pair-classification models. Subsequently, for each under-clustered family, the missing sequences can be reliably predicted using the family-specific alignment score cutoffs obtained in training. We also provide the containerized version of the tool which can be downloaded from https://hub.docker.com/r/akshayayadav/undercl-detection-correction.

